# Profiling RNA-Seq at multiple resolutions markedly increases the number of causal eQTLs in autoimmune disease

**DOI:** 10.1101/128728

**Authors:** Christopher A. Odhams, Deborah S. Cunninghame Graham, Timothy J. Vyse

**Author notes:** Corresponding author (TJV).

## Abstract

Genome-wide association studies have identified hundreds of risk loci for autoimmune disease, yet only a minority (∼25%) share genetic effects with changes to gene expression (eQTLs) in immune cells. RNA-Seq based quantification at whole-gene resolution, where abundance is estimated by culminating expression of all transcripts or exons of the same gene, is likely to account for this observed lack of colocalisation as subtle isoform switches and expression variation in independent exons can be concealed. We performed integrative *cis*-eQTL analysis using association statistics from twenty autoimmune diseases (560 independent loci) and RNA-Seq data from 373 individuals of the Geuvadis cohort profiled at gene-, isoform-, exon-, junction-, and intron-level resolution in lymphoblastoid cell lines. After stringently testing for a shared causal variant using both the Joint Likelihood Mapping and Regulatory Trait Concordance frameworks, we found that gene-level quantification significantly underestimated the number of causal *cis*-eQTLs. Only 5.0-5.3% of loci were found to share a causal *cis*-eQTL at gene-level compared to 12.9-18.4% at exon-level and 9.6-10.5% at junction-level. More than a fifth of autoimmune loci shared an underlying causal variant in a single cell type by combining all five quantification types; a marked increase over current estimates of steady-state causal *cis*-eQTLs. As an example, we dissected in detail the genetic associations of systemic lupus erythematosus and functionally annotated the candidate genes. Many of the known and novel genes were concealed at gene-level (e.g. *BANK1*, *UBE2L3*, *IKZF2*, *TYK2, LYST*). By leveraging RNA-Seq, we were able to isolate the specific transcripts, exons, junctions, and introns modulated by the *cis*-eQTL - which supports the targeted design of follow-up functional studies involving alternative splicing. Causal *cis*-eQTLs detected at different quantification types were also found to localise to discrete epigenetic annotations. We provide our findings from all twenty autoimmune diseases as a web resource.

**Author Summary:** It is well acknowledged that non-coding genetic variants contribute to disease susceptibility through alteration of gene expression levels (known as eQTLs). Identifying the variants that are causal to both disease risk and changes to expression levels has not been easy and we believe this is in part due to how expression is quantified using RNA-Sequencing (RNA-Seq). Whole-gene expression, where abundance is estimated by culminating expression of all transcripts or exons of the same gene, is conventionally used in eQTL analysis. This low resolution may conceal subtle isoform switches and expression variation in independent exons. Using isoform-, exon-, and junction-level quantification can not only point to the candidate genes involved, but also the specific transcripts implicated. We make use of existing RNA-Seq expression data profiled at gene-, isoform-, exon-, junction-, and intron-level, and perform eQTL analysis using association data from twenty autoimmune diseases. We find exon-, and junction-level thoroughly outperform gene-level analysis, and by leveraging all five quantification types, we find >20% of autoimmune loci share a single genetic effect with gene expression. We highlight that existing and new eQTL cohorts using RNA-Seq should profile expression at multiple resolutions to maximise the ability to detect causal eQTLs and candidate genes.

## Introduction

The autoimmune diseases are a family of heritable, often debilitating, complex disorders in which immune dysfunction leads to loss of tolerance to self-antigens and chronic inflammation [1]. Genome-wide association studies (GWAS) have now detected hundreds of susceptibility loci contributing to risk of autoimmunity [2] yet their biological interpretation still remains challenging [3]. Mapping single nucleotide polymorphisms (SNPs) that influence gene expression (eQTLs) can provide meaningful insight into the potential candidate genes and etiological pathways connected to discrete disease phenotypes [4]. For example, such analyses have implicated dysregulation of autophagy in Crohn’s disease [5], the pathogenic role of CD4^+^ effector memory T-cells in rheumatoid arthritis [6], and an overrepresentation of transcription factors in systemic lupus erythematosus [7].

Expression profiling in appropriate cell types and physiological conditions is necessary to capture the pathologically relevant regulatory changes driving disease risk [8]. Lack of such expression data is thought to explain the observed disparity of shared genetic architecture between disease association and gene expression at certain autoimmune loci [9]. A much overlooked cause of this disconnect however, is not only the use of microarrays to profile gene expression, but also the resolution to which expression is quantified using RNA-Sequencing (RNA-Seq) [10]. Expression estimates of whole-genes, individual isoforms and exons, splice-junctions, and introns are obtainable with RNA-Seq [11–18]. The SNPs that affect these discrete units of expression vary strikingly in their proximity to the target gene, localisation to specific epigenetic marks, and effect on translated isoforms [18]. For example, in over 57% of genes with both an eQTL influencing overall gene expression and a transcript ratio QTL (trQTL) affecting the ratio of each transcript to the gene total, the causal variants for each effect are independent and reside in distinct regulatory elements of the genome [18].

RNA-Seq based eQTL investigations that solely rely on whole-gene expression estimates are likely to mask the allelic effects on independent exons and alternatively-spliced isoforms [16–19]. This is in part due to subtle isoform switches and expression variation in exons that cannot be captured at gene-level [20]. A large proportion of trait associated variants are thought to act via direct effects on pre-mRNA splicing that do not change total mRNA levels [21]. Recent evidence also suggests that exon-level based strategies are more sensitive than conventional gene-level approaches, and allow for detection of moderate but systematic changes in gene expression that are not necessarily derived from alternative-splicing events [15,22]. Furthermore, gene-level summary counts can be biased in the direction of extreme exon outliers [22]. Use of isoform-, exon-, and junction-level quantification in eQTL analysis also support the potential to not only point to the candidate genes involved, but also the specific transcripts or functional domains affected [10,18]. This of course facilitates the design of targeted functional studies and better illuminates the causative relationship between regulatory genetic variation and disease. Lastly, though intron-level quantification is not often used in conventional eQTL analysis, it can still provide valuable insight into the role of unannotated exons in reference gene annotations, retained introns, and even intronic enhancers [23,24].

Low-resolution expression profiling with RNA-Seq will impede the subsequent identification of causal eQTLs when applying genetic and epigenetic fine-mapping approaches [25]. In this investigation, we aim to increase our knowledge of the regulatory mechanisms and candidate genes of human autoimmune disease through integration of GWAS and RNA-Seq expression data profiled at gene-, isoform-, exon-, junction-, and intron-level in lymphoblastoid cell lines (LCLs). This is firstly performed in detail using association data from a GWAS in systemic lupus erythematosus, and is then scaled up to a total of twenty autoimmune diseases. Our findings are provided as a web resource to interrogate the functional effects of autoimmune associated SNPs (www.insidegen.com), and will serve as the basis for targeted follow-up investigations.

## Results

### Gene-level expression quantification underestimates the number of causal *cis*-eQTLs

Using densely imputed genetic association data from a large-scale GWAS in systemic lupus erythematosus (SLE) in persons of European descent [7], we performed integrative *cis*-eQTL analysis with RNA-Seq expression data profiled at five resolutions: gene-, transcript-, exon-, junction-, and intron-level. The expression data are derived from the 373 healthy European donors of the Geuvadis project (all individuals are included as part of the 1000 Genomes Project) profiled in lymphoblastoid cell lines (LCLs) [18]. See S1 Figure and methods for a summary of how expression at the five resolutions was quantified using RNA-Seq. A total of 38 genome-wide significant SLE loci (S1 Table) were put forward for analysis following removal of: associated SNPs with minor allele frequency < 5%, secondary associations upon conditional analysis on lead variant, and major histocompatibility complex loci - owing to the known complex linkage disequilibrium (LD) patterns. To test for evidence of a single shared causal variant between disease and gene expression at each of the remaining 38 SLE associated loci, we employed the rigorous Joint Likelihood Mapping (JLIM) framework [9] using summary-level statistics for the SLE association (primary trait) and full genotype-level data for gene expression (secondary trait). Using JLIM, *cis*-eQTLs were defined if a nominal association (*P*<0.01) with at least one SNP existed within 100kb of the SNP most associated with disease and the transcription start site of the gene located within +/-500kb of that SNP (as defined by the authors of the JLIM package). JLIM *P*-values were corrected for multiple testing as per the JLIM standards by using a false discovery rate (FDR) of 5% per RNA-Seq quantification type (i.e. at exon-level, JLIM *P*-values were FDR adjusted for total number of exons tested in *cis* to the 38 SNPs). Causal associations of the integrative *cis*-eQTL SLE GWAS analysis using the JLIM package across the five RNA-Seq quantification types are available in S2 Table and the full output (including non-causal associations) are available in S3 Table. See S2 Figure for the distribution of JLIM *P*-values across the five RNA-Seq quantification types.

We found the number of *cis*-eQTLs driven by the same causal variant as the SLE disease association was markedly underrepresented when considering conventional gene-level quantification (Table 1). Only two of the 38 SLE susceptibility loci (5.3%) were deemed to be causal *cis*-eQTLs at gene-level for three candidate genes. Interestingly, this is a similar proportion to that observed by the authors of the JLIM method (*Chun et al* [9]). They found that 16 of the 272 (5.9%) autoimmune susceptibility loci tested were *cis*-eQTLs driven by a shared causal variant in the Geuvadis RNA-Seq dataset using gene-level quantification (based upon the seven autoimmune diseases interrogated - not including SLE).

**Table 1.**
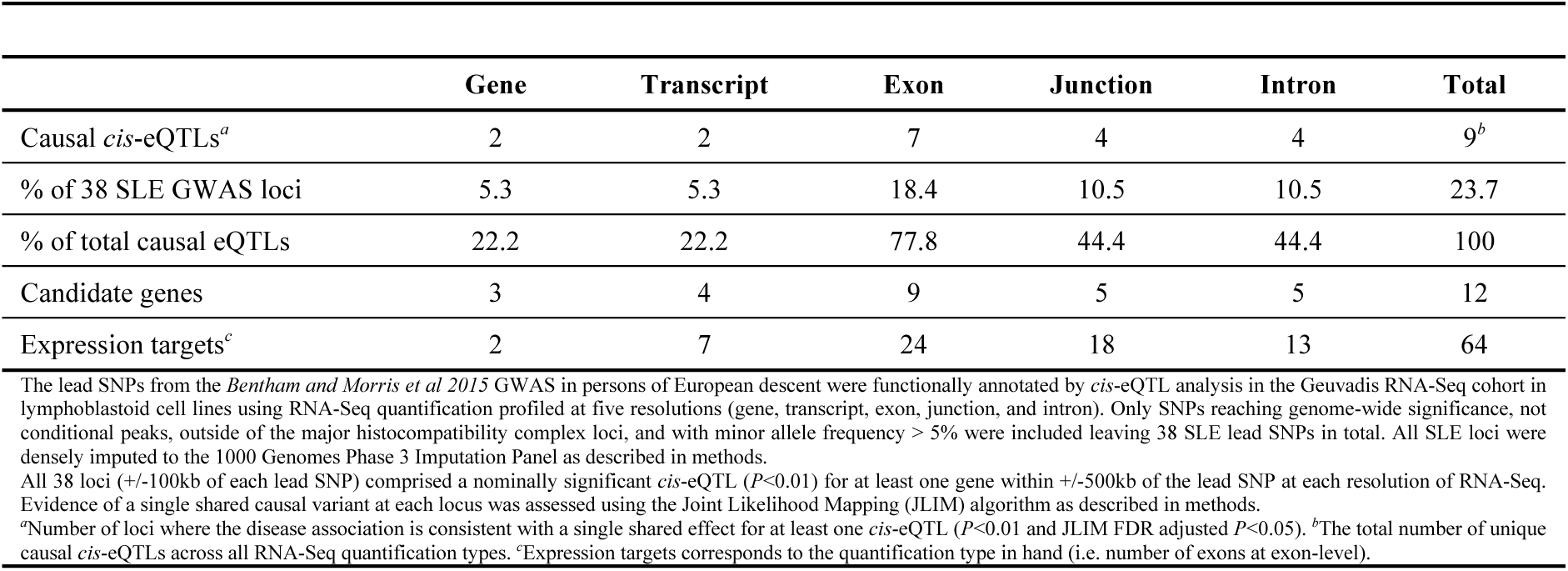
Number of *cis*-eQTLs driven by the same causal variant as the SLE disease association (total number of SLE loci: 38)

|  | Gene | Transcript | Exon | Junction | Intron | Total |
| --- | --- | --- | --- | --- | --- | --- |
| Causal *cis*-eQTLs<sup>a</sup> | 2 | 2 | 7 | 4 | 4 | 9<sup>b</sup> |
| % of 38 SLE GWAS loci | 5.3 | 5.3 | 18.4 | 10.5 | 10.5 | 23.7 |
| % of total causal eQTLs | 22.2 | 22.2 | 77.8 | 44.4 | 44.4 | 100 |
| Candidate genes | 3 | 4 | 9 | 5 | 5 | 12 |
| Expression targets<sup>c</sup> | 2 | 7 | 24 | 18 | 13 | 64 |

The lead SNPs from the *Bentham and Morris et al 2015* GWAS in persons of European descent were functionally annotated by *cis*-eQTL analysis in the Geuvadis RNA-Seq cohort in lymphoblastoid cell lines using RNA-Seq quantification profiled at five resolutions (gene, transcript, exon, junction, and intron). Only SNPs reaching genome-wide significance, not conditional peaks, outside of the major histocompatibility complex loci, and with minor allele frequency > 5% were included leaving 38 SLE lead SNPs in total. All SLE loci were densely imputed to the 1000 Genomes Phase 3 Imputation Panel as described in methods.
All 38 loci (+/-100kb of each lead SNP) comprised a nominally significant *cis*-eQTL ( $P<0.01$ ) for at least one gene within +/-500kb of the lead SNP at each resolution of RNA-Seq. Evidence of a single shared causal variant at each locus was assessed using the Joint Likelihood Mapping (JLIM) algorithm as described in methods.
<sup>a</sup>Number of loci where the disease association is consistent with a single shared effect for at least one *cis*-eQTL ( $P<0.01$ and JLIM FDR adjusted $P<0.05$ ). <sup>b</sup>The total number of unique causal *cis*-eQTLs across all RNA-Seq quantification types. <sup>c</sup>Expression targets corresponds to the quantification type in hand (i.e. number of exons at exon-level).

Of note, transcript-level quantification did not increase the number of causal *cis*-eQTLs (Table 1). Transcript-level analysis did, however, yield a greater number of candidate genes (seven individual transcripts derived from a total of four genes). Both junction- and intron-level quantification increased the number of causal *cis*-eQTLs to four (10.5% of the 38 total SLE loci). Using exon-level quantification, we were able to define seven of the 38 SLE susceptibility loci (18.4%) as being significant *cis*-eQTLs driven by a single shared causal variant. Exon-level analysis also produced the greatest number of candidate gene targets: nine unique genes derived from 24 individual SNP-exon pairs (Table 1). Therefore, even with multiple testing burden to correct for all SNP-exon *cis*-eQTL pairs; we firstly conclude that exon-level analysis detects more causal *cis*-eQTLs than gene-level.

### A fifth of associated SNPs possess shared genetic effects with *cis*-eQTLs using RNA-Seq in LCLs

By combining all five types of RNA-Seq quantification (gene, transcript, exon, junction, and intron) we could define nine of the 38 SLE susceptibility loci (23.7%) as being driven by the same causal variant as the *cis*-eQTL in LCLs (Table 1). Interestingly, this value, derived from interrogating only a single cell type, is almost equal to the total number of causal autoimmune *cis*-eQTLs detected by *Chun et al* [9] (∼25%) when looking across the three different cell types analysed using JLIM (CD4^+^ T-cells – measured by microarray, CD14^+^ monocytes – microarray, and LCLs – RNA-Seq gene-level).

We found that when considering the specificity of *cis*-eQTLs and target genes identified by JLIM mapping across the five RNA-Seq quantification types, both gene- and transcript-level quantification were redundant with respect to exon-level data; i.e. there were no causal *cis*-eQTLs or target genes detected at gene- or transcript-level that were not captured by exon-level analysis (S3 Figure). Both junction- and intron-level quantification captured a single causal *cis*-eQTL each that was not captured by exon-level. We conclude that profiling at all resolutions of RNA-Seq is required to capture the full set of potentially causal *cis*-eQTLs.

### Associated SNPs are most likely to colocalize with exon- and junction-level *cis*-eQTLs

We compared the detection of *cis*-eQTLs using a standard linear-regression approach with the JLIM method. To fully explore relationships within our results, a pairwise comparison was made across the five RNA-Seq quantification types for matched SNP-gene *cis*-eQTL pairs (Figure 1). We only considered matched SNP-gene *cis*-eQTL association pairs that had a nominal *cis*-eQTL association *P*-value < 0.01 in both quantification types, and to be conservative, when multiple transcripts, exons, junctions, and introns were annotated with the same gene symbol, we selected the associations that minimized the difference in JLIM *P*-value between matched SNP-gene *cis*-eQTLs across RNA-Seq quantification types. There were over 250 matched SNP-gene *cis*-eQTL pairs per comparison. We firstly observed that the correlation of both *cis*-eQTL association *P*-values from regression and JLIM *P*-values across RNA-Seq quantification types reflected the methods in which expression quantification was obtained (Figure 1A). Both *cis*-eQTL and JLIM *P*-values between matched SNP-gene pairs at gene- and transcript-level were highly correlated as gene-level estimates are obtained from the sum of all transcript-level estimates for the same gene (see methods and S1 Figure). Exon-level and junction-level associations were also highly correlated due to split-reads being incorporated into the exon-level estimate. As expected, intron-level *cis*-eQTL and JLIM *P*-values for matched SNP-gene pairs were only weakly correlated against other quantification types - as reads mapping to introns are not included in the other quantification models. Interestingly, although *cis*-eQTL association *P*-values for matched SNP-gene pairs between transcript-level and junction-level were found to be relatively high (*r*^2^=0.70), we found the JLIM *P*-values for the matched pairs to be comparatively low (*r*^2^=0.29); suggesting that whilst the strength of the *cis*-eQTL maybe similar between these quantification types, the underlying causal variants driving the disease and *cis*-eQTL association are likely to be independent.

**Figure 1.**
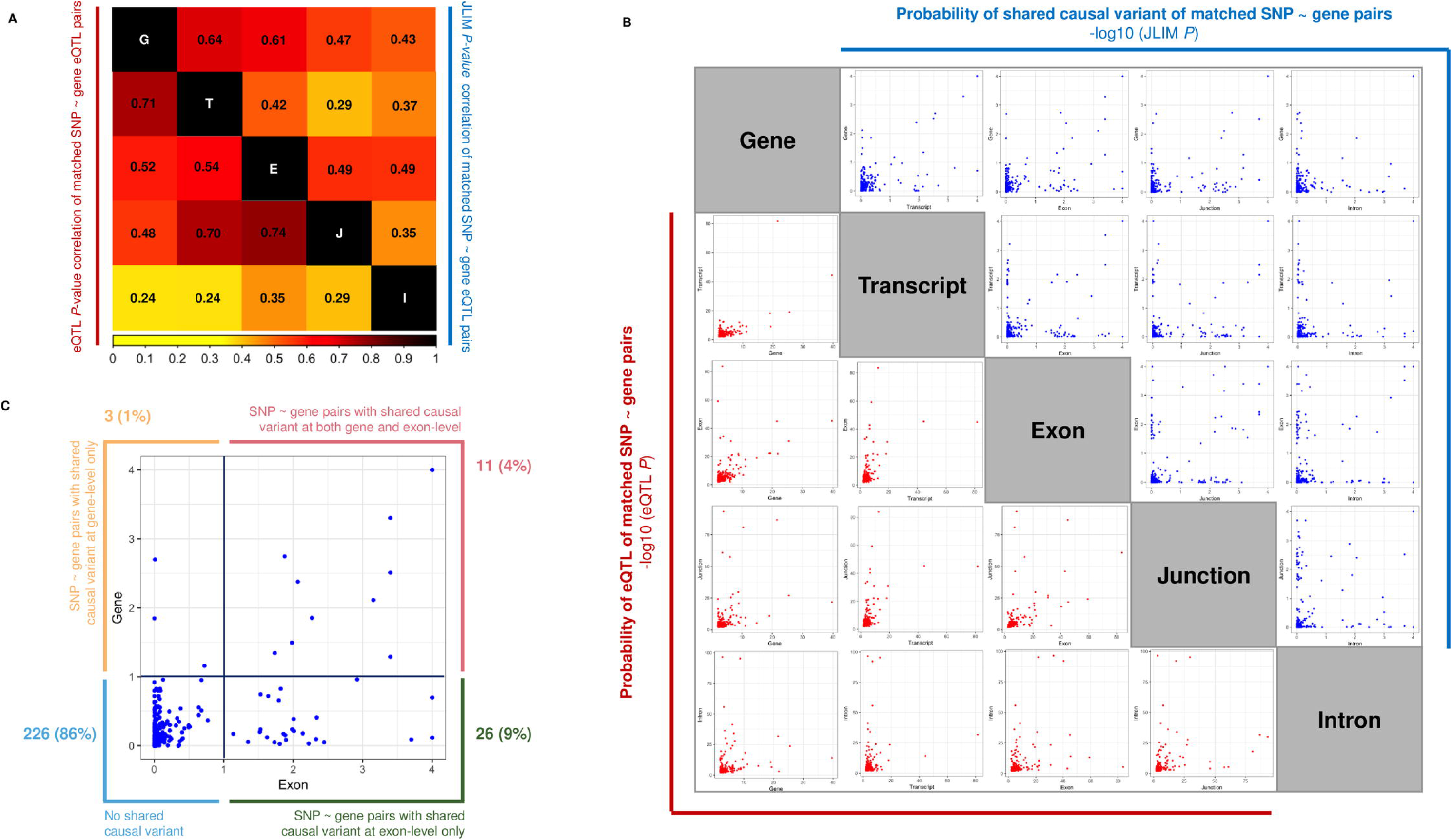
Pairwise comparison of *cis*-eQTL and JLIM *P*-values for matched SNP-gene pairs. This figure is complementary to the data in Table 2 and is derived from *cis*-eQTL analysis of the 38 SLE associated SNPs using RNA-Seq and implementation of the JLIM method to assess evidence of a shared causal variant. (A) We measured the Pearson’s correlation separately of all *cis*-eQTL and JLIM *P*-values between matched SNP-gene *cis*-eQTL pairs across the five RNA-Seq quantification types. We only considered matched SNP-gene *cis*-eQTL association pairs that had a nominal *cis*-eQTL association *P*-value < 0.01 in both quantification types, and to be conservative, when multiple transcripts, exons, junctions, and introns were annotated with the same gene symbol, we selected the associations that minimized the difference in JLIM *P*-value between matched SNP-gene *cis*-eQTLs across RNA-Seq quantification types. Note the weak JLIM *P*-value correlation of matched transcript-level and junction-level *cis*-eQTLs suggesting they stem from independent causal variants. (B) Correlation plots of matches SNP-gene *cis*-eQTL pairs as described above (red: *cis*-eQTL *P*-value; blue: JLIM *P*-value). Note that JLIM *P*-values often aggregate on the axis rather than on the diagonal suggesting independent causal variants across different quantification types. (C) An example of the sensitivity of exon-level analysis relative to gene-level. The majority of nominally significant JLIM *P*-values (<0.01) for matched SNP-gene pairs are captured by exon-level analysis and concealed at gene-level (green box: 9%).

By plotting the JLIM *P*-values for matched SNP-gene pairs between different quantification types, we found many instances of *P*-values distributed along the axes rather than on the diagonal (Figure 1B). Our findings therefore suggest that often, one quantification type is more likely to explain the observed disease association than the other. When we compared conventional gene-level *cis*-eQTL analysis against exon-level results (Figure 1C), we found that of the 296 matched SNP-gene *cis*-eQTL associations (*P*<0.01), eleven (4%) were deemed to share the same causal variant at both gene- and exon-level using a nominal JLIM *P*-value threshold < 0.01. Only three of the 296 matched SNP-gene *cis*-eQTL associations (1%) were captured by gene-level only - in contrast to the 26 (9% of total associations) captured uniquely at exon-level. As expected, the overwhelming majority of *cis*-eQTL associations (86%) did not possess a single shared causal variant at either gene- or exon-level. We performed this analysis for all possible combinations of quantification types (Table 2). In each instance, gene-level analysis detected only the minority of nominally causal associations for matched SNP-gene association pairs (JLIM *P*<0.01). Exon-level and junction-level analysis consistently detected more causal *cis*-eQTL associations than gene-, transcript-, and intron-level. In fact, when combined, exon- and junction-level analysis explained the most nominally causal associations for all significant SNP-gene *cis*-eQTL association pairs (23.8%).

**Table 2.**
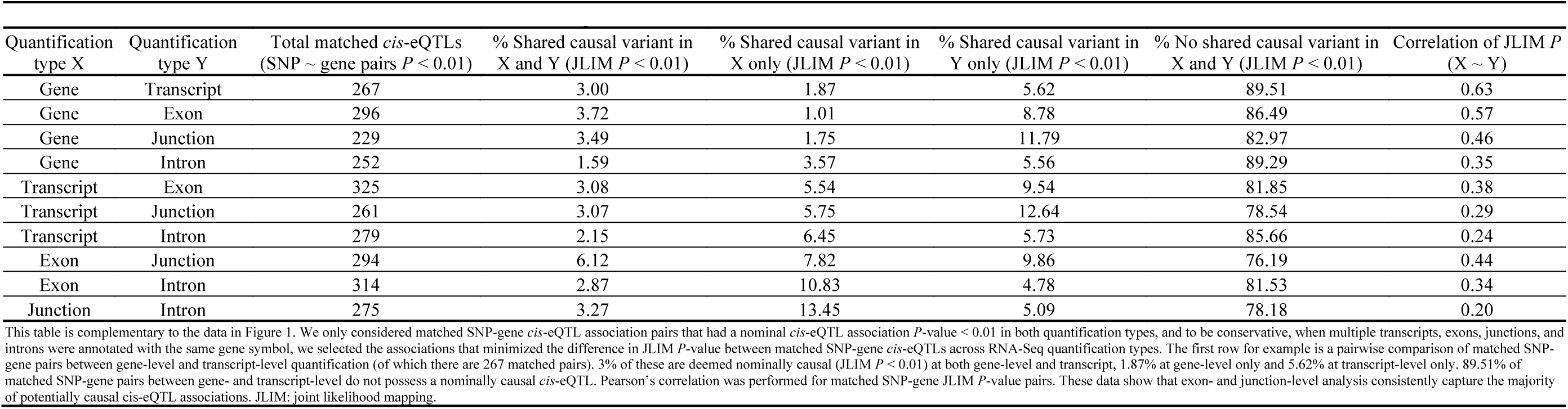
Pairwise comparison of the number of *cis*-eQTLs with a nominal JLIM *P*-value < 0.01.

|  |  |  |  |  |  |  |  |
| --- | --- | --- | --- | --- | --- | --- | --- |
| Quantification type X | Quantification type Y | Total matched *cis*-eQTLs (SNP ~ gene pairs *P* < 0.01) | % Shared causal variant in X and Y (JLIM *P* < 0.01) | % Shared causal variant in X only (JLIM *P* < 0.01) | % Shared causal variant in Y only (JLIM *P* < 0.01) | % No shared causal variant in X and Y (JLIM *P* < 0.01) | Correlation of JLIM *P* (X ~ Y) |
| Gene | Transcript | 267 | 3.00 | 1.87 | 5.62 | 89.51 | 0.63 |
| Gene | Exon | 296 | 3.72 | 1.01 | 8.78 | 86.49 | 0.57 |
| Gene | Junction | 229 | 3.49 | 1.75 | 11.79 | 82.97 | 0.46 |
| Gene | Intron | 252 | 1.59 | 3.57 | 5.56 | 89.29 | 0.35 |
| Transcript | Exon | 325 | 3.08 | 5.54 | 9.54 | 81.85 | 0.38 |
| Transcript | Junction | 261 | 3.07 | 5.75 | 12.64 | 78.54 | 0.29 |
| Transcript | Intron | 279 | 2.15 | 6.45 | 5.73 | 85.66 | 0.24 |
| Exon | Junction | 294 | 6.12 | 7.82 | 9.86 | 76.19 | 0.44 |
| Exon | Intron | 314 | 2.87 | 10.83 | 4.78 | 81.53 | 0.34 |
| Junction | Intron | 275 | 3.27 | 13.45 | 5.09 | 78.18 | 0.20 |

### Leveraging RNA-Seq aids GWAS interpretation and reveals novel candidate genes

We functionally dissected the 12 candidate genes taken from the nine SLE associated loci that showed strong evidence of a shared causal variant with a *cis*-eQTL in LCLs. The nine, causal *cis*-eQTLs and corresponding 12 candidate genes per RNA-Seq quantification type are listed in Table 3 along with their *cis*-eQTL association *P*-values and related JLIM *P*-values. We systematically annotated all 12 genes using a combination of cell/tissue expression patterns, mouse models, known molecular phenotypes, molecular interactions, and associations with other autoimmune diseases (S4 Table). We found the majority of novel SLE candidate genes detected by RNA-Seq were predominately expressed in immune-related tissues such as whole blood, the spleen and thymus, and the small intestine. Based on our gene annotation and what is already documented at certain loci, we were sceptical on the pathogenic involvement of three candidate genes (*PHTF1*, *ARHGAP30*, and *RABEP1*). Although the *cis*-eQTL effect for these genes is evidently driven by the shared causal variant as the disease association (defined by JLIM), it is possible that these effects of expression modulation are merely passengers that are carried on the same functional haplotype as the true causal gene(s) and do not contribute themselves to the breakdown of self-tolerance (detailed in S4 Table). We show the regional association plots and the candidate genes detected from *cis*-eQTL analysis in S4 Figure.

**Table 3.** Nine SLE loci contain *cis*-eQTLs driven by the same variant as the disease association.

|  |  |  |  |  |  |  |  |  |  |  |  |
| --- | --- | --- | --- | --- | --- | --- | --- | --- | --- | --- | --- |
|  |  | Gene |  | Transcript |  | Exon |  | Junction |  | Intron |  |
| Lead SNP | Gene | eQTL *P*<sup>a</sup> | JLIM *P* | eQTL *P* | JLIM *P* | eQTL *P* | JLIM *P* | eQTL *P* | JLIM *P* | eQTL *P* | JLIM *P* |
| rs2476601 | *PHTF1* | - | - | 2.2 x 10<sup>-3</sup> | 6.2 x 10<sup>-1</sup> | 5.0 x 10<sup>-8</sup> | 1 | 8.4 x 10<sup>-47</sup> | 1 | **1.4 x 10<sup>-4</sup>** | **1.0 x 10<sup>-4</sup>** |
| rs1801274 | *ARHGAP30* | 2.4 x 10<sup>-6</sup> | 8.1 x 10<sup>-1</sup> | - | - | **1.1 x 10<sup>-4</sup>** | **2.0 x 10<sup>-4</sup>** | 9.4 x 10<sup>-3</sup> | 7.4 x 10<sup>-3</sup> | 1.2 x 10<sup>-3</sup> | 4.8 x 10<sup>-1</sup> |
| rs9782955 | *LYST* | 5.4 x 10<sup>-3</sup> | 3.90 x 10<sup>-1</sup> | 8.0 x 10<sup>-6</sup> | 9.8 x 10<sup>-1</sup> | 1.6 x 10<sup>-3</sup> | 4.6 x 10<sup>-3</sup> | **1.3 x 10<sup>-3</sup>** | **2.0 x 10<sup>-4</sup>** | 1.0 x 10<sup>-5</sup> | 5.0 x 10<sup>-1</sup> |
| rs3768792 | *IKZF2* | - | - | 1.5 x 10<sup>-3</sup> | 7.7 x 10<sup>-1</sup> | **1.9 x 10<sup>-4</sup>** | **3.0 x 10<sup>-4</sup>** | 1.0 x 10<sup>-5</sup> | 9.0 x 10<sup>-1</sup> | **1.1 x 10<sup>-5</sup>** | **2.0 x 10<sup>-4</sup>** |
| rs10028805 | *BANK1* | 1.8 x 10<sup>-3</sup> | 3.1 x 10<sup>-3</sup> | 4.9 x 10<sup>-3</sup> | 3.2 x 10<sup>-3</sup> | **1.8 x 10<sup>-5</sup>** | **4.0 x 10<sup>-4</sup>** | **2.5 x 10<sup>-4</sup>** | **2.0 x 10<sup>-4</sup>** | 1.8 x 10<sup>-4</sup> | 9.7 x 10<sup>-1</sup> |
| rs2736340 | *BLK* | **3.2 x 10<sup>-26</sup>** | < 10<sup>-4</sup> | **1.0 x 10<sup>-9</sup>** | < 10<sup>-4</sup> | **1.4 x 10<sup>-31</sup>** | < 10<sup>-4</sup> | **7.6 x 10<sup>-28</sup>** | < 10<sup>-4</sup> | **3.1 x 10<sup>-24</sup>** | < 10<sup>-4</sup> |
|  | *FAM167A* | **2.3 x 10<sup>-40</sup>** | < 10<sup>-4</sup> | **4.4 x 10<sup>-45</sup>** | < 10<sup>-4</sup> | **5.1 x 10<sup>-46</sup>** | < 10<sup>-4</sup> | **1.5 x 10<sup>-22</sup>** | < 10<sup>-4</sup> | **7.4 x 10<sup>-15</sup>** | < 10<sup>-4</sup> |
| rs2286672 | *RABEP1* | 1.4 x 10<sup>-3</sup> | 5.1 x 10<sup>-2</sup> | 1.3 x 10<sup>-4</sup> | 9.4 x 10<sup>-1</sup> | **7.4 x 10<sup>-5</sup>** | **4.0 x 10<sup>-4</sup>** | 4.5 x 10<sup>-4</sup> | 7.0 x 10<sup>-4</sup> | 1.3 x 10<sup>-4</sup> | 8.5 x 10<sup>-1</sup> |
| rs2304256 | *TYK2* | 1.2 x 10<sup>-3</sup> | 7.6 x 10<sup>-1</sup> | 9.9 x 10<sup>-6</sup> | 9.9 x 10<sup>-1</sup> | **2.5 x 10<sup>-9</sup>** | < 10<sup>-4</sup> | 1.3 x 10<sup>-4</sup> | 3.0 x 10<sup>-3</sup> | **2.2 x 10<sup>-9</sup>** | **2.0 x 10<sup>-4</sup>** |
|  | *ATG4D* | - | - | 3.8 x 10<sup>-3</sup> | 7.2 x 10<sup>-3</sup> | 6.4 x 10<sup>-5</sup> | 3.8 x 10<sup>-3</sup> | **3.8 x 10<sup>-4</sup>** | **2.0 x 10<sup>-4</sup>** | 6.6 x 10<sup>-5</sup> | 9.7 x 10<sup>-1</sup> |
| rs7444 | *UBE2L3* | 5.7 x 10<sup>-3</sup> | 2.0 x 10<sup>-1</sup> | **5.9 x 10<sup>-14</sup>** | < 10<sup>-4</sup> | **9.9 x 10<sup>-5</sup>** | < 10<sup>-4</sup> | 5.1 x 10<sup>-5</sup> | 9.5 x 10<sup>-1</sup> | 1.2 x 10<sup>-3</sup> | 9.0 x 10<sup>-1</sup> |
|  | *CCDC116* | **2.5 x 10<sup>-5</sup>** | **5.0 x 10<sup>-4</sup>** | 1.4 x 10<sup>-6</sup> | 3.0 x 10<sup>-4</sup> | **4.9 x 10<sup>-4</sup>** | **4.0 x 10<sup>-4</sup>** | - | - | - | - |
| Nine of the 38 SLE loci (24%) were found to be driven by the same causal variant as the disease association across all five RNA-Seq quantification types in LCLs (*cis*-eQTL *P*<0.01 and joint likelihood of shared association FDR<0.05). Bold type indicates associations that show evidence of a shared causal variant for *cis*-eQTL and disease. <sup>a</sup>Minimum *cis*-eQTL *P*-value for any SNP within 100 kb of the lead SNP. Dashes (-) indicate genes that were either not detected or had minimum *cis*-eQTL *P*>0.01 in the RNA-Seq quantification type in hand. JLIM *P*-values <10<sup>-4</sup> indicates the JLIM statistic is more extreme than permutation. JLIM: joint likelihood mapping. If multiple SNP-unit associations are deemed to be causal (i.e. one SNP shows a causal association to two exons of the same gene, the association with the smallest JLIM *P*-value is reported). |  |  |  |  |  |  |  |  |  |  |  |

The causal *cis*-eQTL rs2736340 for genes *BLK* and *FAM167A* was detected at all RNA-Seq profiling types. It is well established that the risk allele of this SNP reduces proximal promoter activity of *BLK*; a member of the Src family kinases that functions in intracellular signalling and the regulation of B-cell proliferation, differentiation, and tolerance [26]. The allelic consequence of *FAM167A* expression modulation is unknown. We found multiple instances of known SLE susceptibility genes that were concealed when using gene-level quantification. For example, we defined rs7444 as a causal *cis*-eQTL for *UBE2L3* at transcript- and exon-level - but not at gene-level (Table 3). The risk allele of rs7444 has been associated with increased expression of *UBE3L3* (Ubiquitin conjugating enzyme E2 L3) in *ex vivo* B-cells and monocytes and correlates with NF-κB activation along with increased circulating plasmablast and plasma cell numbers [27]. Similarly, the rs10028805 SNP is a known splicing *cis*-eQTL for *BANK1* (B-cell scaffold protein with ankyrin repeats 1). We replicated at exon-, and junction-level this splicing effect which has been proposed to alter the B-cell activation threshold [28]. Again, this mechanism was not detected using gene-level quantification.

*IKZF2* (detected at the exon-level only) is a transcription factor thought to play a key role in T-reg stabilisation in the presence of inflammatory responses [29]. *IKZF2* deficient mice acquire an auto-inflammatory phenotype in later life similar to rheumatoid arthritis, with increased numbers of activated CD4^+^ and CD8^+^ T-cells, T-follicular helper cells, and germinal centre B-cells, which culminates in autoantibody production [30]. Of note, other members of this gene family, *IKZF1* and *IKZF3*, are also associated with SLE and can hetero-dimerize (S4 Table) [7]. We also believe *LYST*, *ATG4D*, and *TYK2* to also be intriguing candidate genes. *LYST* encodes a lysosomal trafficking regulator [31] whilst *ATG4D* is a cysteine peptidase involved in autophagy and this locus is associated with multiple sclerosis, psoriasis, and rheumatoid arthritis [32]. *TYK2* is discussed in greater detail in the following section.

### RNA-Seq can resolve the potential causal regulatory mechanism(s)

Interestingly, for the three causal SNP-gene pairs detected at gene-level (rs2736340 – *BLK*, rs2736340 – *FAM167A*, and rs7444 – *CCDC116*), we found that at exon-level, all expressed exons of the stated genes were deemed to possess causal associations. For example, rs2736340 is a causal *cis*-eQTL for all thirteen exons of *BLK* and for all three exons of *FAM167A* (S5 Table). These data suggest that gene-level analysis is capturing associations where all - or the majority of exons - are modulated by the *cis*-eQTL in a causal manner.

We found that within the SLE associated loci that showed evidence of a shared causal variant with a *cis*-eQTL (Table 3), there were many instances in which the proposed causal *cis*-eQTL modulated expression of only a single expression element. This enabled us to resolve the potential regulatory effect of the causal *cis*-eQTL to a particular transcript, exon, junction, or intron (S5 Table). We were able to resolve to a single expression element in nine of the twelve candidate SNP-gene pairs. For example, rs9782955 is a causal *cis*-eQTL for *LYST* at junction-level for only a single junction (chr1:235915471-235916344; *cis*-eQTL *P*=1.3x10^−03^; JLIM *P*=2.0x10^−04^). We provide depicted examples of this isolation analysis for candidate genes *IKZF2* (S5 Figure), *UBE2L3* (S6 Figure), and *LYST* (S7 Figure). Clearly when only the minority of exons are effected – which we found occurred in nine of twelve association pairs - gene-level analysis conceals the *cis*-eQTL association.

We provide a worked example of resolving the causal mechanism(s) using RNA-Seq for the novel association rs2304256 with *TYK2* (Figure 2). The top panel of Figure 2A shows the genetic association to SLE at the 19p13.2 susceptibility locus tagged by lead SNP rs2304256 (*P*=1.54x10^−12^). Multiple tightly correlated SNPs span the gene body and the 3′ region of *TYK2* – which encodes Tyrosine Kinase 2 - thought to be involved in the initiation of type I IFN signalling [33]. In the panel below, we plot the gene-level association of all SNPs in *cis* to *TYK2* and show no significant association of rs3204256 with *TYK2* expression (*P*=0.18). At exon-, and intron-level, we were able to classify rs2304256 as a causal *cis*-eQTL for a single exon (chr19: 10475527-10475724; *cis*-eQTL *P*=2.58x10^−09^; JLIM *P*<10^−04^) and single intron (chr19: 10473333-10475290; *P*=2.20x10^−08^; JLIM *P*=2x10^−04^) of *TYK2* respectively as shown in the bottom two panels of Figure 2A. We show the exon and intron labelling of *TYK2* in further detail in S8 Fig. We found strong correlation of association *P*-values of the SLE GWAS and the *P*-values of *TYK2 cis*-eQTLs against at exon-level and intron-level, but not at gene-level; strengthening our observation that rs2304256 is a causal *cis*-eQTL for *TYK2* at these resolutions (Figure 2B). The risk allele rs2304256 [C] was found to be associated with decreased expression of the *TYK2* exon and increased expression of the *TYK2* intron (Figure 2C). By plotting the *cis*-eQTL *P*-values alongside the JLIM *P*-values for all exons and introns of *TYK2* against rs2304256 (Figure 2D), we clearly show that only a single exon and a single intron of *TYK2* colocalize with the SLE association signal – marked by an asterisk (note that rs2304256 is a strong *cis*-eQTL for many introns of *TYK2* but only shares a causal variant with one intron). We show the genomic location of the affected exon and intron of *TYK2* in Figure 2E (exon 8 and the intron between exons 9 and 10 – N.B that exons and introns are numbered based on their inclusion in the *cis*-eQTL analysis and some maybe omitted from analysis due to no expression). Intron 9-10 of *TYK2* is clearly ‘expressed’ in LCLs according to transcription levels assayed by RNA-Seq on LCLs (GM12878) from ENCODE (Figure 2E).

**Figure 2.**
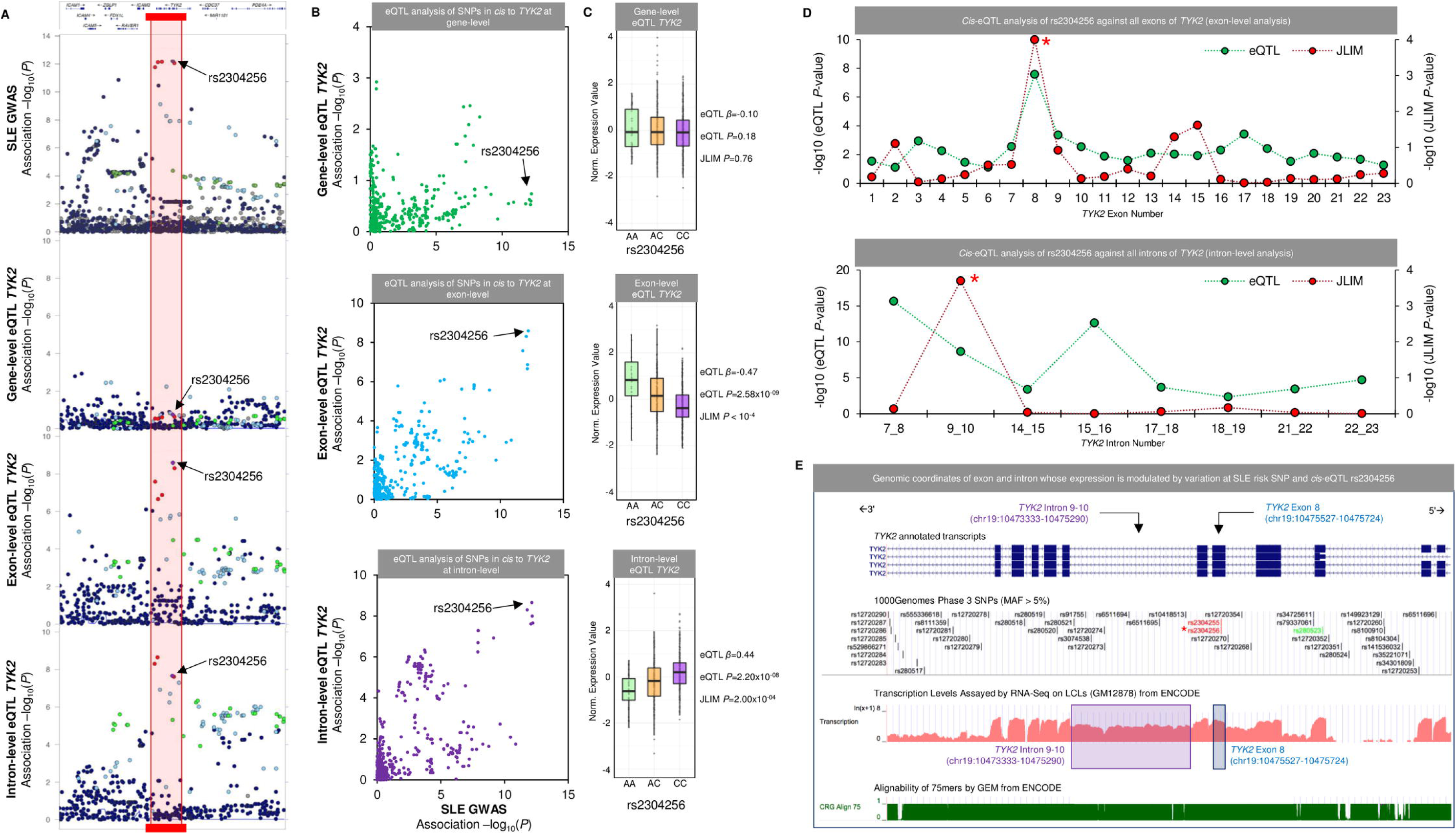
Isolation of potential causal molecular mechanism in *TYK2* by SLE *cis*-eQTL rs2304256. (A) SLE GWAS association plot and *cis*-eQTL association plot around the 19p13.2 susceptibility locus tagged by rs2304256. The top panel shows the association plot with SLE that spans the gene body and 3′ region of *TYK2* (Tyrosine Kinase 2). The haplotype block composed of highly correlated SNPs is highlighted in the red block. The second panel shows the *cis*-eQTL association plot at gene-level of all proximal SNPs to *TYK2* (no significant association with rs2304256 is detected). The third panel shows the same regional association but at exon-level for the most associated exon of *TYK2* with rs2304256 – the bottom panel is at intron-level for *TYK2* (both are highly associated). (B) Correlation of SLE GWAS *P*-value and *cis*-eQTL association *P*-value for all SNPs in *cis* to *TYK2*. We show at gene-level the most associated SLE SNPs are not *cis*-eQTLs (top panel). The middle and bottom panels show the same correlation at exon-level and intron-level and reveal the most associated SNPs to SLE are also the most associated *cis*-eQTLs to *TYK2*. (C) The direction of effect of *cis*-eQTL rs2304256 with *TYK2* at gene-level (top), exon-level (middle), and intron-level (bottom panel). The risk allele is rs2304256 [C]. (D) The top panel shows *cis*-eQTL association and JLIM *P*-values for all exons of *TYK2* against rs2304256. Exon 8 (marked by an asterisk) is defined as having a causal association with rs2304256. The bottom panel shows the intron-level *cis*-eQTL of *TYK2* against rs2304256. Note many introns are *cis*-eQTLs but are not causal with rs2304256. Exons and introns are numbered consecutively from start to end of gene if they are expressed (note some are not and therefore not included). (E) The genomic location of the single exon and single intron of *TYK2* that are modulated by rs2304256 are highlighted (rs2304256 is marked by an asterisk in red). The bottom two panels show the transcription levels assayed by RNA-Seq on LCLs assayed by ENCODE. Note intron 9-10 of *TYK2* is clearly expressed. The alignability of 75-mers by GEM is also shown to show the mapability of reads around rs2304256.

Interestingly, rs2304256 (marked by an asterisk in Figure 2E) is a missense variant (V362F) within the affected exon 8 of *TYK2*. The PolyPhen prediction of this substitution is predicted to be benign and, to the best of our knowledge, no investigation has isolated the functional effect of this particular amino acid change. We do not believe the *cis*-eQTL at exon 8 to be a result of variation at rs3204256 and mapping biases, as the alignability of 75mers by GEM from ENCODE is predicted to be robust around exon 8 (Figure 2E). In fact, rs3204256 [C] is the reference allele yet is associated with decreased expression of exon 8.

In conclusion, we have found an interesting and novel mechanism that would have been concealed by gene-level analysis that involves the risk allele of a missense SNP associated with decreased expression of a single exon of *TYK2* but increased expression of the neighbouring intron. Whether the *cis*-eQTL effect and missense variation act in a combinatorial manner and whether the intron is truly retained or if it is derived from an unannotated transcript of *TYK2* is an interesting line of investigation.

### Detection of *cis*-eQTLs and candidate-genes of autoimmune disease using RNA-Seq

We re-performed our integrative *cis*-eQTL analysis with the same Geuvadis RNA-Seq dataset in LCLs using association data from twenty autoimmune diseases. This was to firstly reiterate the importance of leveraging RNA-Seq in GWAS interpretation and to secondly demonstrate that our findings in SLE persisted across other immunological traits. As the raw genetic association data were not available for all twenty diseases, we were unable to implement the JLIM pipeline which requires densely typed or imputed GWAS summary-level statistics. We therefore opted to use the Regulatory Trait Concordance (RTC) method, which requires full genotype-level data for the expression trait, but only the marker identifier for the lead SNP of the disease association trait (see methods for a description of the RTC method). We stringently controlled our integrative *cis*-eQTL analysis for multiple testing to limit potential false positive findings of overlapping association signals. To do this, we applied a Bonferroni correction to nominal *cis*-eQTL *P*-values separately per disease and per RNA-Seq quantification type (i.e. at exon-level, *cis*-eQTL *P*-values were corrected for the total number of exons tested in *cis* the associated SNPs of the single disease in hand). A similar strategy was adopted by the authors of the JLIM package who corrected separately for specific disease and cell type combinations [9]. We rigorously defined causal *cis*-eQTLs, as associations with *P*BF < 0.05 and RTC ≥ 0.95. An overview of the analysis pipeline is depicted in S9 Figure and S10 Figure. Using an *r*^2^ cut-off of 0.8 and a 100kb limit, we pruned the 752 associated SNPs from the twenty human autoimmune diseases from the Immunobase resource (S6 Table) to obtain 560 independent susceptibility loci. Again, we only considered common (MAF >5%), autosomal loci outside of the MHC.

Our findings confirmed our previous results from the SLE investigation and again support the gene-level study using the JLIM package from *Chun et al* [9]. As before, we found that only 5% (28 of the 560 loci) of autoimmune susceptibility loci were deemed to share causal variants with *cis*-eQTLs using either gene- or transcript-level analysis (Figure 3A). Exon-level analysis more than doubled the yield to 13% (72 of the 560 loci) with junction-, and intron-level analysis also outperforming gene-level (10% and 8% respectively). When combining all RNA-Seq quantification types, we could define 20% of autoimmune associated loci (110 of the 560 loci) as being candidate causal *cis*-eQTLs - which corroborates our previous estimate in SLE using the JLIM package (23.7%).

**Figure 3.**
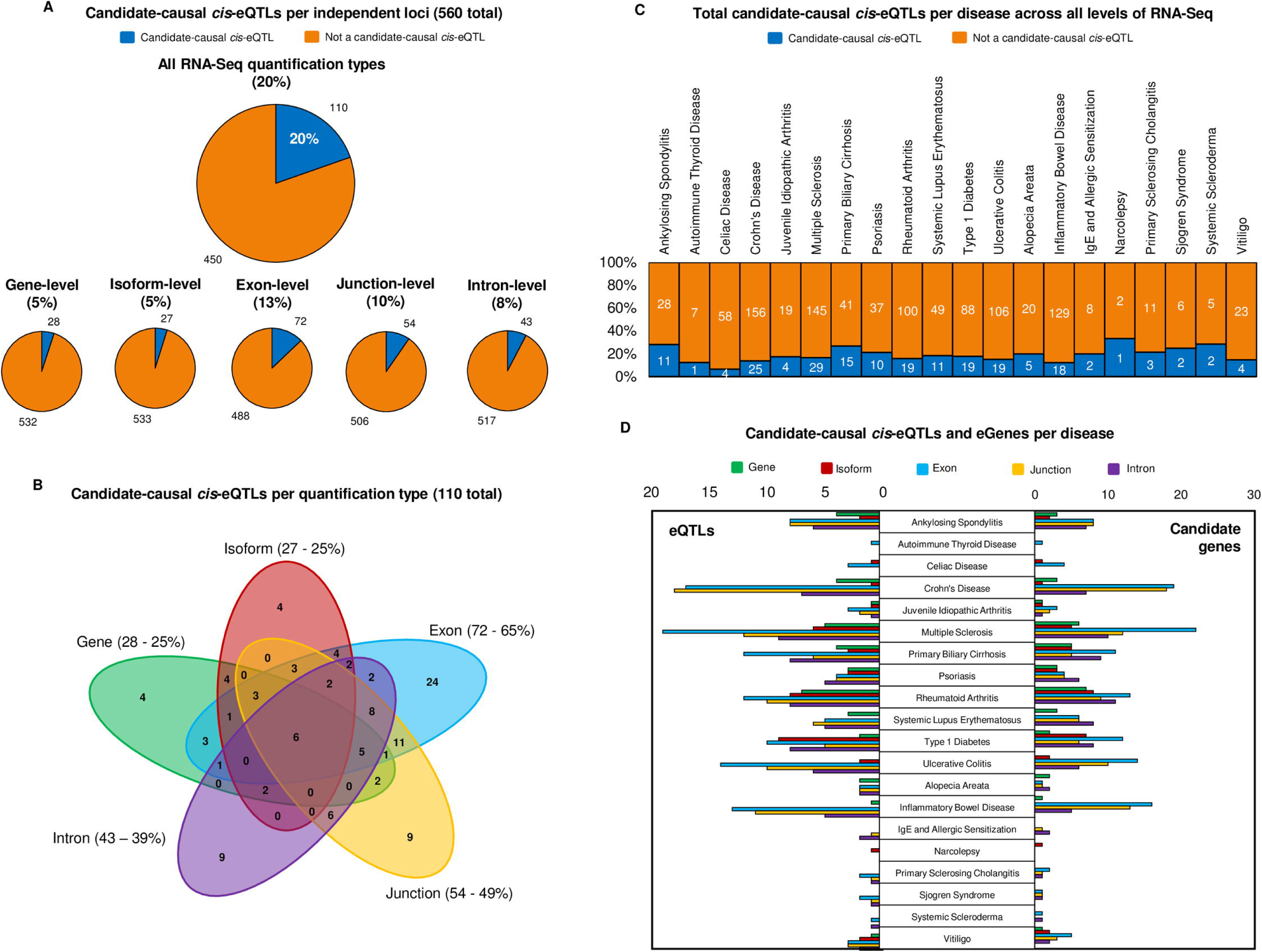
Breakdown of autoimmune associated causal *cis*-eQTLs using RNA-Seq. (A) Percentage and number of causal *cis*-eQTL associations detected per RNA-Seq quantification type, following LD pruning of associated SNPs from twenty autoimmune diseases to 560 independent susceptibly loci. The top chart shows the number of causal *cis*-eQTLs when combining all RNA-Seq profiling types together (20%). (B) Sharing of causal *cis*-eQTL associations per quantification type (110 detected in total). Percentage of causal *cis*-eQTLs captured are shown as a percentage of the 110 total. (C) Total causal *cis*-eQTLs per disease across all five levels of RNA-Seq quantification, using the 20 diseases of the ImmunoBase resource. In orange are disease-associated SNPs that show no shared association with expression across any quantification type. In blue are the disease-associated SNPs that are also causal *cis*-eQTLs. (D) Causal *cis*-eQTLs and candidate genes per disease broken down by quantification type.

By separating causal *cis*-eQTL associations out by quantification type, we found over half (65%) were detected at exon-level, and considerable overlap of *cis*-eQTL associations existed between both types (Figure 3B). Unlike in our SLE analysis, gene- and isoform-level analysis did capture a small fraction of causal *cis*-eQTLs that were not captured at exon-level. Our data therefore suggest that although exon- and junction-level, and to a lesser extent intron-level analysis, capture most candidate-causal *cis*-eQTLs. It is necessary to prolife gene-expression at all quantification types to avoid misinterpretation of the functional impact of disease associated SNPs.

We mapped the causal *cis*-eQTLs detected by all RNA-Seq quantification types back to the diseases to which they are associated (Figure 3C). Interestingly, we observed the diseases that fell below the 20% average comprised autoimmune disorders related to the gut: celiac disease (7%), inflammatory bowel disease (14%), Crohn’s disease (16%), and ulcerative colitis (18%). These observations are likely to be a result of the cellular expression specificity of associated genes in colonic tissue and in T-cells [34]. Correspondingly, we observed an above-average frequency of causal *cis*-eQTLs detected in SLE (22%) and primary biliary cirrhosis (37%); diseases in which the pathogenic role of B-lymphocytes and autoantibody production is well documented [34]. Note that there are 60 SLE GWAS associations in this analysis as these originate from three independent GWA studies (S6 Table). We further broke down our results per disease by RNA-Seq quantification type (Figure 3D) and in all cases, the greatest frequency of causal *cis*-eQTLs and candidate genes were captured by exon- and junction-level analyses.

### Web resource for functional interpretation of association studies of autoimmune disease

We provide our analysis as a web resource (found at www.insidegen.com) for researchers to lookup causal *cis*-eQTLs and candidate genes from the twenty autoimmune diseases detected across the five RNA-Seq quantification types. The data are sub-settable and exportable by SNP ID, gene, RNA-Seq resolution, genomic position, and association to specific autoimmune diseases.

### Causal *cis*-eQTLs localise to discrete chromatin regulatory elements

The causal variants underling *cis*-eQTL associations at the five RNA-Seq quantification types were often independent (Figure 1) and a previous investigation has suggested that causal variants of gene-level and transcript-level *cis*-eQTLs reside in discrete functional elements of the genome [18]. We therefore investigated whether this notion held true across the five RNA-Seq quantification types tested in this study. To accomplish this, we selected the causal *cis*-eQTLs from the twenty autoimmune diseases interrogated, and per quantification type, tested for enrichment of these SNPs across various chromatin regulatory elements taken from the Roadmap Epigenomics Project in LCLs (using both the Roadmap chromatin state model and the positions of histone modifications). We implemented the permutation-based GoShifter algorithm to test for enrichment of causal *cis*-eQTLs and tightly correlated variants (*r*^2^>0.8) in genomic functional annotations in LCLs (see methods) [25]. Results of this analysis are depicted in Figure 4. We found the 28 gene-level *cis*-eQTLs were enriched in two chromatin marks: strong enhancers (*P*=0.036) and H3K27ac occupancy sites – a marker of active enhancers (*P*=0.002). Transcript-level *cis*-eQTLs were also enriched in H3K27ac occupancy sites (*P*=0.039) but were not enriched in any other marks. The 72 exon-level *cis*-eQTLs were additionally enriched in active promoters (*P*=0.017). Interestingly, the 54 causal *cis*-eQTLs detected at junction-level were found to be enriched in weak enhancers only (*P*=0.002); whilst the 43 intron-level *cis*-eQTLs were enriched in chromatin states predicted to be involved in transcriptional elongation (*P*=0.001; 83% of intron-level *cis*-eQTLs). Disease relevant *cis*-eQTLs detected at different expression phenotypes using RNA-Seq clearly localise to largely discrete functional elements of the genome.

**Figure 4.**
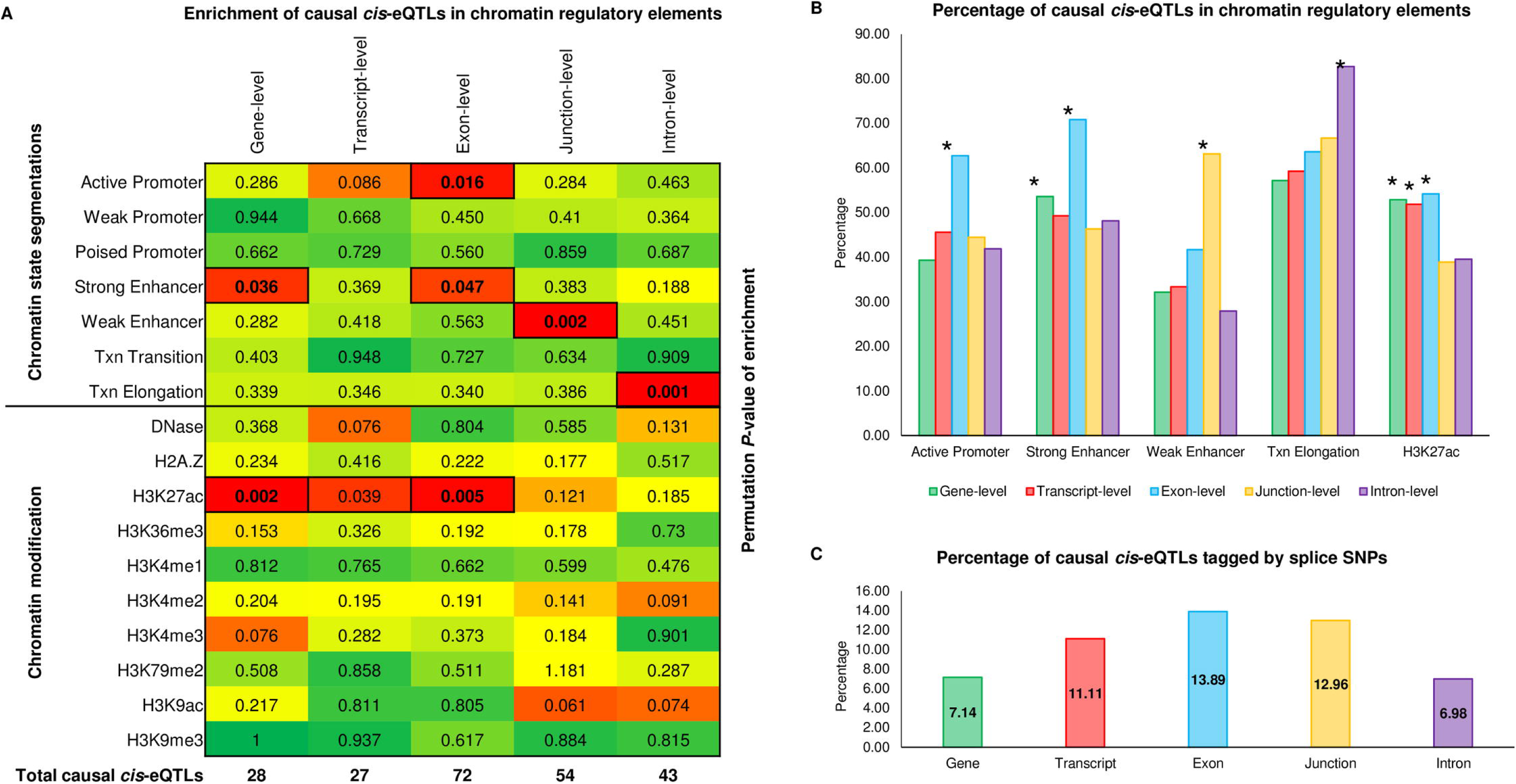
Functional annotation of causal autoimmune *cis*-eQTLs. (A) We took the causal autoimmune *cis*-eQTLs detected for each RNA-Seq quantification type and performed enrichment testing for chromatin state segmentation and histone marks in LCLs taken from the NIH Roadmap Epigenomics Project. We used the GoShifter algorithm to do this (see methods); which takes all SNPs in strong LD (*r*2>0.8) with the causal *cis*-eQTLs and calculates the proportion of SNPs overlapping chromatin marks, the positions of the marks are then shuffled whilst retaining the SNP positions, and the fraction of overlap recalculated over 1,000 permutations. A permutation *P*-value is then generated – which is annotated in each box (*P*<0.05 deemed significant). The heat colour is representative of the permutation *P*-value. Significant enrichment tests are highlighted in bold. The total number of causal *cis*-eQTLs per quantification type are annotated at the bottom of the heatmap. (B) The percentage of causal *cis*-eQTLs in chromatin regulatory marks per quantification type. An asterisk shows that this level of enrichment is deemed to be significant as shown in panel A. (C) The percentage of causal *cis*-eQTLs in chromatin regulatory marks per quantification type that are or are highly correlated (*r*2>0.8) with SNPs that alter splice site consensus sequences of the target genes (assessed by Sequence Ontology for the hg19 GENCODE v12 reference annotation).

We quantified the number of causal *cis*-eQTLs and tightly correlated variants (*r*^2^>0.8) per quantification type that were predicted to be alter splice site consensus sequences of the target genes (assessed by Sequence Ontology for the hg19 GENCODE v12 reference annotation). We found only two of the 28 (7%) gene-level *cis*-eQTLs disrupted consensus splice-sites for their target genes compared to the 14% and 13% detected at exon- and junction-level respectively (Figure 4C). Our data suggest that although exon- and junction- level analysis leads to the greatest frequency of causal *cis*-eQTLs, the majority at this resolution cannot be explained directly by variation in annotated splice site consensus sequences (splice region/donor/acceptor/ variants).

## Discussion

Elucidation of the functional consequences of non-coding genetic variation in human disease is a major objective of medical genomics [35]. Integrative studies that map disease-associated eQTLs in relevant cell types and physiological conditions are proving essential in progression towards this goal through identification of causal SNPs, candidate-genes, and illumination of molecular mechanisms [36]. In autoimmune disease, where there is considerable overlap of immunopathology, integrative eQTL investigations have been able to connect discrete aetiological pathways, cell types, and epigenetic modifications, to particular clinical manifestations [2,34,36,37]. Emerging evidence however has suggested that only a minority (∼25%) of autoimmune associated SNPs share casual variants with basal-level *cis*-eQTLs in primary immune cell-types [9].

Genetic variation can influence expression at every stage of the gene regulatory cascade - from chromatin dynamics, to RNA folding, stability, and splicing, and protein translation [21]. It is now well documented that SNPs affecting these units of expression vary strikingly in their genomic positions and localisation to specific epigenetic marks [18]. The eQTLs that affect pre-transcriptional regulation - affecting all isoforms of a gene - differ in the proximity to the target gene and effect on translated isoforms than their co-transcriptional trQTL (transcript ratio QTL) counterparts. Where the effect size of eQTLs generally increases in relation to transcription start site proximity, trQTLs are distributed across the transcript body and generally localise to intronic binding sites of splicing factors [18,21]. In over 57% of genes with both an eQTL influencing overall gene expression and an trQTL affecting the ratio of each transcript to the gene total, the causal variants for each effect are independent and reside in distinct regulatory elements of the genome [18]. In fact, three primary molecular mechanisms are thought to link common genetic variants to complex traits. A large proportion of trait associated SNPs act via direct effects on pre-mRNA splicing that do not change total mRNA levels [21]. Common variants also act via alteration of pre-mRNA splicing indirectly through effects on chromatin dynamics and accessibility. Such chromatin accessibility QTLs are however more likely to alter total mRNA levels than splicing ratios. Lastly, it is thought that only a minority of trait associated variants have direct effects on total gene expression that cannot be explained by changes in chromatin. As RNA-Seq becomes the convention for genome-wide transcriptomics, it is essential to maximise its ability to resolve and quantify discrete transcriptomic features so to expose the genetic variants that contribute to changes in expression and isoform usage. The reasoning for our investigation therefore was to delineate the limits of microarray and RNA-Seq based eQTL cohorts in the functional annotation of autoimmune disease association signals.

To map autoimmune disease associated *cis*-eQTLs, we interrogated RNA-Seq expression data profiled at gene-, isoform, exon-, junction-, and intron-level, and tested for a shared genetic effect at each significant association. As we had densely imputed summary statistics from our SLE GWAS, we opted to use the Joint Likelihood Mapping (JLIM) framework [9] to test for a shared causal variant between the disease and *cis*-eQTL signals. This framework has been rigorously benchmarked against other colocalisation procedures. Summary statistics were not available for the remaining autoimmune diseases and therefore we implemented the Regulatory Trait Concordance (RTC) method for these diseases and set a stringent multiple testing threshold to define causal *cis*-eQTLs. We found the estimates of causal *cis*-eQTLs were near identical between the two methods used (Table 1 and Figure 3A). Exon- and junction-level quantification led to the greatest frequency of causal *cis*-eQTLs and candidate genes (exon-level: 13-18%, junction-level: JLIM: 10-11%). We conclusively found that associated variants were in fact more likely to colocalize with exon- and junction-level *cis*-eQTLs when applying a nominal JLIM *P*-value threshold of <0.01 (Figure 1B and Table 2). Gene-level analysis was thoroughly outperformed in all cases (5%). Our findings that gene-level analysis explain only 5% of causal *cis*-eQTLs corroborate the findings from *Chun et al* [9] who composed and used the JLIM framework to annotate variants associated with seven autoimmune diseases (multiple sclerosis, IBD, Crohn’s disease, ulcerative colitis, T1D, rheumatoid arthritis, and celiac disease). They found that only 16 of the 272 autoimmune associated loci (6%) shared causal variants with *cis*-eQTLs using gene-level RNA-Seq (with the same Geuvadis European cohort in LCLs as used herein). In our investigation, we argue that it is necessary to profile expression at all possible resolutions to diminish the likelihood of overlooking potentially causal *cis*-eQTLs. In fact, by combining our results across all resolutions, we found that 20-24% of autoimmune loci were candidate-causal *cis*-eQTLs for at least one target gene. Our study therefore increases the number of autoimmune loci with shared genetic effects with *cis*-eQTLs in a single cell type by over four-fold. Interestingly, using microarray data from CD4^+^ T-cells *Chun et al* classified 37 of the 272 autoimmune loci (14%) as causal *cis*-eQTLs [9] - strengthening the hypothesis that autoimmune loci (especially those associated with inflammatory diseases of the gut) are enriched in CD4^+^ T-cell subsets and the cells themselves are pathogenic [25,34]. Microarray data are known to underestimate the number of true causal *cis*-eQTLs [10]. If we assume that by leveraging RNA-Seq we can increase the number of causal *cis*-eQTLs four-fold, we hypothesise that as many as ∼54% of autoimmune loci may share causal *cis*-eQTLs with gene expression at multiple resolutions in CD4^+^ T-cell populations. A large RNA-Seq based eQTL cohort profiled across many CD4^+^ T-cell subsets will therefore be of great use when annotating autoimmune-related traits. We reason that although using relevant cell types and context-specific conditions will undoubtedly increase our understanding of how associated variants alter cell physiology and ultimately contribute to disease risk; it is clearly shown herein that we are only picking the low hanging fruit in current eQTL analyses. We argue it necessary to reanalyse existing RNA-Seq based eQTL cohorts at multiple resolutions and ensure new datasets are similarly dissected. Despite the severe multiple testing burden, we also argue that expression profiling at multiple resolutions using RNA-Seq may be advantageous even when looking for *trans*-eQTL effects. As *trans*-eQTLs are generally more cell-type specific and have a weaker effect size, we decided not to perform such analyses using the Geuvadis LCL data. Large RNA-Seq based eQTL cohorts in whole-blood will be more suitable for such analysis [19].

As well as biological reasons for using multiple expression phenotypes for integrative eQTL analysis, there are also technical factors to consider. Gene-level expression estimates can generally be obtained in two ways – union-exon based approaches [14,17] and transcript-based approaches [11,12]. In the former, all overlapping exons of the same gene are merged into union exons, and intersecting exon and junction reads (including split-reads) are counted to these pseudo-gene boundaries. Using this counting-based approach, it is also possible to quantify meta-exons and junctions easily and with high confidence by preparing the reference annotation appropriately [13,15,38]. Introns can be quantified in a similar manner by inverting the reference annotation between exons and introns [18]. Of note, we found intron-level quantification generated more candidate-causal *cis*-eQTLs than gene-level (Figure 3A). As the library was synthesised from poly-A selection, these associations are unlikely due to differences in pre-mRNA abundance. Rather, they are likely derived from either true retained introns in the mature RNA or from coding exons that are not documented in the reference annotation used. Transcript-based approaches make use of statistical models and expectation maximization algorithms to distribute reads among gene isoforms - resulting in isoform expression estimates [11,12]. These estimates can then be summed to obtain the entire expression estimate of the gene. Greater biological insight is gained from isoform-level analysis; however, disambiguation of specific transcripts is not trivial due to substantial sequence commonality of exons and junctions. In fact, we found only 5% of autoimmune loci shared a causal variant at transcript-level.

The different approaches used to estimate expression can also lead to significant differences in the reported counts. Union-based approaches, whilst computationally less expensive, can underestimate expression levels relative to transcript-based, and this difference becomes more pronounced when the number of isoforms of a gene increases, and when expression is primarily derived from shorter isoforms [20]. The Geuvadis study implemented a transcript-based approach to obtain whole-gene expression estimates. Clearly therefore, a gold standard of reference annotation and eQTL mapping using RNA-Seq is essential for comparative analysis across datasets. Our findings support recent evidence that suggests exon-level based strategies are more sensitive and specific than conventional gene-level approaches [22]. Subtle isoform variation and expression of less abundant isoforms are likely to be masked by gene-level analysis. Exon-level allows for detection of moderate but systematic changes in gene expression that are not captured at gene-level, and also, gene-level summary counts can be shifted in the direction of extreme exon outliers [22]. It is therefore important to note that a positive exon-level eQTL association does not necessarily mean a differential exon-usage or splicing mechanism is involved; rather a systematic expression effect across the whole gene may exist that is only captured by the increased sensitivity. Additionally, by combining exon-level with other RNA-Seq quantification types, inferences can be made on the particular isoforms and functional domains affected by the eQTL which can later aid biological interpretation and targeted follow-up investigations [10]. We clearly show this from our analysis of SLE candidate genes *IKZF2* (S5 Figure), *UBE2L3* (S6 Figure), *LYST* (S7 Figure) and *TYK2* (Figure 2). For *TYK2* we reveal a novel mechanism whereby the associated variant rs2304256 [C] leads to decreased expression of a single exon and increased expression of a neighbouring intron (Figure 2). By isolating particular exons, junctions, and introns, one can design more refined follow-up investigations to study the functional impact of non-coding disease associated variants. We show how our findings can be leveraged to comprehensively examine GWAS results of autoimmune diseases. We found nine of the 38 SLE susceptibility loci were causal *cis*-eQTLs (Table 3) for 12 candidate genes which we later functionally annotated in detail (S4 Table).

Taken together, we have provided a deeper mechanistic understanding of the genetic regulation of gene expression in autoimmune disease by profiling the transcriptome at multiple resolutions using RNA-Seq. Similar analyses leveraging RNA-Seq in new and existing datasets using relevant cell types and context-specific conditions (such as response eQTLs as shown in [39]) will undoubtedly increase our understanding of how associated variants alter cell physiology and ultimately contribute to disease risk.

## Materials and Methods

### RNA-Sequencing expression data in lymphoblastoid cell lines

RNA-Sequencing (RNA-Seq) expression data from 373 lymphoblastoid cell lines (LCLs) derived from four European sub-populations (Utah Residents with Northern and Western European Ancestry, British in England and Scotland, Finnish in Finland, and Toscani in Italia) of the Geuvadis project [18] were obtained from the EBI ArrayExpress website under accession: E-GEUV-1. The 89 individuals of the Geuvadis project from the Yoruba in Ibadan, Nigeria were excluded from this analysis. All individuals were included as part of the 1000Genomes Project. Expression was profiled using RNA-Seq at five quantification types: gene-, transcript-, exon-, junction-, and intron-level (the files downloaded and used in this analysis have the suffix: ‘QuantCount.45N.50FN.samplename.resk10.txt.gz’). Full methods of expression quantification can be found in the original publication and on the Geuvadis wiki page: http://geuvadiswiki.crg.es/). We have also provided a breakdown of the quantification methods in S1 Figure. Expression data downloaded represent quantifications that are corrected for sequencing depth and gene/exon etc length (RPKM). Only expression elements quantified in >50 % of individuals were kept and Probabilistic Estimation of Expression Residuals (PEER) had been used to remove technical variation [40]. We transformed all expression data to a standard normal distribution.

In summary, transcripts, splice-junctions, and introns were quantified using Flux Capacitor against the GENCODE v12 basic reference annotation [16]. Reads belonging to single transcripts were predicted by deconvolution per observations of paired-reads mapping across all exonic segments of a locus. Gene-level expression was calculated as the sum of all transcripts per gene. Annotated splice junctions were quantified using split read information, counting the number of reads supporting a given junction. Intronic regions that are not retained in any mature annotated transcript, and reported mapped reads in different bins across the intron to distinguish reads stemming from retained introns from those produced by not yet annotated exons. Meta-exons were quantified by merging all overlapping exonic portions of a gene into non-redundant units and counting reads within these bins. Reads were excluded when the read pairs map to two different genes.

### SLE associated SNPs

SNPs genetically associated to systemic lupus erythematosus (SLE) were taken from the *Bentham and Morris et al 2015* GWAS in persons of European descent [7]. The study comprised a primary GWAS, with validation through meta-analysis and replication study in an external cohort (7,219 cases, 15,991 controls in total). Independently associated susceptibility loci taken forward for this investigation were those that passed either genome-wide significance (*P*<5x10^−08^) in the primary GWAS or meta-analysis and/or those that reached significance in the replication study (q<0.01). We defined the lead SNP at each locus as either being the SNP with the lowest *P*-value post meta-analysis or the SNP with the greatest evidence of a missense effect as defined by a Bayes Factor (see original publication). We omitted non-autosomal associations and those within the Major Histocompatibility Complex (MHC), and SNPs with a minor allele frequency (MAF) < 0.05. In total, 38 independently associated SLE associated GWAS SNPs were taken forward for investigation (S1 Table). Each susceptibility locus had previously been imputed to the level of 1000 Genomes Phase3 using a combination of pre-phasing by the SHAPEIT algorithm and imputation by IMPUTE (see original publication for full details) [7].

### *Cis*-eQTL analysis and Joint Likelihood Mapping (JLIM) of SLE associated SNPs

#### Primary trait summary statistics file

A JLIM index file for each of the 38 SLE associated SNPs was firstly generated by taking the position of each SNP (hg19) and a creating a 100kb interval in both directions. Summary-level association statistics were obtained form the *Bentham and Morris et al* 2015 European SLE GWAS (imputed to 1000Genomes Phase 3). We downloaded summary-level association data (chromosome, position, SNP, *P*-value) for all directly typed or imputed SNPs with an IMPUTE info score ≥0.7 within each of the 38 intervals. The two-sided *P*-value was transformed into a *Z*-statistic as described by JLIM.

#### Reference LD file

Genotype files in VCF format for all 373 European individuals of the Geuvadis RNA-Seq project were obtained from the EBI ArrayExpress under accession: E-GEUV-1. The 41 individuals genotyped on the Omni 2.5M SNP array had been previously imputed to the Phase 1 v3 release as described [18]; the remaining had been sequenced as part of the 1000 Genomes Phase1 v3 release (low-coverage whole genome and high-coverage exome sequencing data). Using VCFtools, we created PLINK binary ped/map files for each of the 38 intervals and kept only biallelic SNPs with a MAF >0.05, imputation call-rates ≥ 0.7, Hardy–Weinberg equilibrium *P*-value >1x10^‒04^ and SNPs with no missing genotypes, we also only included SNPs that we had primary trait association summary statistics for. These are referred to as the secondary trait genotype files. We then used the JLIM Perl script *fetch.refld0.EUR.pl* to generate the 38 reference LD files from the 373 individuals (the script had been edited to include the extra 95 Finnish individuals).

#### Cis-eQTL analysis

We created a separate PLINK phenotype file (sample ID, normalized expression residual) for each individual gene, transcript, exon, junction, and intron in *cis* (within +/-500kb) to the 38 lead SLE GWAS SNPs. We only included protein-coding, lincRNA, and antisense genes in our analysis as classified by Ensembl BioMart. Using the chromosome 20 genotype VCF file of the 373 European individuals (E-GEUV-1), we conducted principle component analysis (PCA) and generated an identity-by-state matrix using the Bioconductor package SNPRelate (S9 Figure) [41]. Based on these results, we decided to include the first three principle components and the binary imputation status (as 41 individuals had been genotyped on the Omni 2.5M SNP array were imputed to the Phase 1 v3 release) of the European individuals (derived from Phase1 and Phase2 1000Genomes releases) in the *cis*-eQTL analysis so to minimize biases derived from population structure and imputation status.

We used PLINK to perform *cis*-eQTL analysis using the ‘*--linear*’ function, including the above covariates, for each expression unit (phenotype file) in *cis* to the 38 loci (secondary trait genotype files). We performed 10,000 permutations per regression and saved the output of each permutation procedure. In *cis* to the 38 SLE SNPs were: 439 genes, 1,448 transcripts (originating from 456 genes), 3,045 exons (400 genes), 2,886 junctions (332 genes), and 1,855 introns (443 genes).

#### Joint likelihood mapping (JLIM) and multiple testing correction

Per RNA-Seq quantification type, a JLIM configuration file was created using the *jlim_gencfg.sh* script and JLIM then run using *run_jlim.sh* – setting the *r*^2^ resolution limit to 0.8. We merged the configuration files and output files to create the final results table which included the primary and secondary trait association *P*-value, the JLIM statistic, and the JLIM *P*-value by permutation. Multiple testing was corrected for on the JLIM *P*-values per RNA-Seq quantification type using a false discovery rate (FDR) as applied by the authors of JLIM. A JLIM *P*-value <10^−04^ means that the JLIM statistic is more extreme than the permutation (10,000). We classified causal *cis*-eQTLs as SLE associated variants that share a single causal variant with a *cis*-eQTL based on the following: if there existed a nominal *cis*-eQTL (*P*<0.01) with at least one SNP within 100kb of the SNP most associated with disease, the transcription start site of the expression target was located within +/-500kb of that SNP, and the FDR adjusted JLIM *P*-value of the association passed the 5% threshold. Candidate genes modulated by the causal *cis*-eQTL.

### Functional annotation of SLE associated genes from *cis*-eQTL analysis

Using publically available resources, we systematically annotated the twelve SLE associated genes that were classified as being modulated by causal *cis*-eQTLs. The expression profiles at RNA-level across multiple cell and tissue types were interrogated in GTEx [42] and the Human Protein Atlas [43] - with the top three cell/tissue types documented per gene. We noted using Online Mendelian Inheritance in Man [44] any gene-phenotype relationships by caused by allelic variants and any immune-related phenotypes of animal models. Protein-protein interactions of candidate genes were taken from the BioPlex v2.0 interaction network (conducted in HEK293T cells) [45]. Using the ImmunoBase resource (https://www.immunobase.org/), we looked up each gene and noted if the gene had been prioritized as the ‘candidate gene’ within the susceptibility locus per publication. Finally, we counted the number publications from PubMed found using the keywords ‘gene name AND SLE’.

### Associated SNPs from twenty autoimmune diseases

Autoimmune associated SNPs were taken from the ImmunoBase resource (www.immunobase.org). This resource comprises summary case-control association statistics from twenty diseases: twelve originally targeted by the ImmunoChip consortium (ankylosing spondylitis, autoimmune thyroid disease, celiac disease, Crohn’s disease, juvenile idiopathic arthritis, multiple sclerosis, primary biliary cirrhosis, psoriasis, rheumatoid arthritis, systemic lupus erythematosus, type 1 diabetes, ulcerative colitis), and eight others (alopecia areata, inflammatory bowel disease, IgE and allergic sensitization, narcolepsy, primary sclerosing cholangitis, Sjogren syndrome, systemic scleroderma, vitiligo).

The curated studies and their corresponding references used in this analysis are presented in S6 Table. For each disease, we took the lead SNPs which were defined as a genome-wide significant SNP with the lowest reported *P*-value in a locus. Associations on the X-chromosome and within the MHC and SNPs with minor allele frequency < 5% were omitted from analysis, leaving 752 associated SNPs. We pruned these loci using the ‘*--indep-pairwise*’ function of PLINK 1.9 with a window size of 100kb and an *r*^2^ threshold of 0.8, to create an independent subset of 560 loci.

### Integrative *cis*-eQTL analysis of twenty autoimmune diseases with RNA-Seq

An overview of the integration pipeline using the twenty autoimmune diseases against the Geuvadis RNA-Seq cohort in 373 European LCLs is depicted in S10 Figure. Genotype data of the 373 individuals were transformed and quality controlled as previously described in the above methods sections (biallelic SNPs kept with a MAF >0.05, imputation call-rates ≥ 0.7, Hardy–Weinberg equilibrium *P*-value >1x10^−04^).

We opted to use the Regulatory Trait Concordance (RTC) method to assess the likelihood of a shared causal variant between the disease association and the *cis*-eQTL signal [46]. This method requires full genotype-level data for the expression trait but only the marker identifier for the lead SNP of the disease association trait. SNPs within the 560 associated loci for the expression trait were firstly classified according to their position in relation to recombination hotspots (based on genome-wide estimates of hotspot intervals) [47]. Normalized gene expression residuals (PEER factor normalized RPKM) for each quantification type were transformed to standard normal and the first three principle components used as covariates in the *cis*-eQTL model as well as the binary imputation status (as previously described above). All *cis*-eQTL association testing was performed using a liner regression model in R. *Cis*-eQTL mapping was performed for the lead SNP and all SNPs within the hotspot recombination interval against protein-coding, lincRNA, and antisense expression elements (genes, transcripts, exons etc.) within +/-500kb of the lead SNP. In *cis* to the 560 loci were: 7,633 genes, 27,257 transcripts (originating from 7,310 genes), 52,651 exons (5,435 genes), 48,627 junctions (4,237 genes), 34,946 introns (6,233 genes).

For each *cis*-eQTL association, the residuals from the linear-regression of the best *cis*-asQTL (lowest association *P*-value within the hotspot interval) were extracted. Linear regression was then performed using all SNPs within the defined hotspot interval against these residuals. The RTC score was then calculated as (*N_SNPs_*-*Rank_GWAS SNP_ / N_SNPs_*). Where *N_SNPs_* is the total number of SNPs in the recombination hotspot interval, and *Rank_GWAS SNP_* is the rank of the GWAS SNP association *P*-value against all other SNPs in the interval from the liner association against the residuals of the best *cis*-eQTL.

We rigorously adjusted for multiple testing of *cis*-eQTL *P*-values using a Bonferroni correction per quantification type (corrected for number of genes, isoforms, exons, junctions, and introns tested) and per disease – as we wanted to keep our analysis as close to the authors of JLIM who themselves also adjusted per cell type and per disease. We stringently defined causal *cis*-eQTLs as associations with expression *P*_BF_ < 0.05 and an RTC score ≥ 0.95. Candidate genes are modulated by the *cis*-eQTL.

### Functional enrichment of causal *cis*-eQTLs in chromatin regulatory elements

To test for enrichment of causal *cis*-eQTL associations in chromatin regulatory elements we implemented the Genomic Annotation Shifter (GoShifter) package [25]. Chromatin regulatory elements were divided into two categories: chromatin state segmentation and histone marks. The genomic coordinates of the fifteen predicted chromatin state segmentations (active promoter, strong enhancer, insulator etc.) for LCLs (in the GM12878 cell-line) were downloaded from the UCSC Table browser (track name: wgEncodeBroadHmmGm12878HMM). Histone marks and DNase hypersensitivity sites were obtained from the NIH Roadmap Epigenomics Project for LCLs (GM12878) in NarrowPeak format. Sites were filtered for genome-wide significance using an FDR threshold of 0.01 and peak widths harmonised to 200bp in length centred on the peak summit (as used in the GoShifter publication).

We obtained all SNPs in strong LD (*r*^2^> 0.8) with the causal *cis*-eQTLs by using the *getLD.sh* script from GoShifter (interrogating the 1000Genomes Project for Phase3 Europeans). Per quantification type, we then calculated the proportion of loci in which at least one SNP in LD overlapped a chromatin regulatory element (conducted one at a time per chromatin mark). The coordinates of the chromatin marks were then randomly shifted, whilst retaining the positions of the SNPs, and frequency of overlap re-calculated. This was carried out over 1,000 permutations to draw the null distribution. The *P*-value was calculated as the proportion of iterations for which the number of overlapping loci was equal to or greater than that for the tested SNPs (*P* < 0.05 used as significance threshold).

### Data visualisation and online resource

R version 3.3.1 and ggplot2 was used to create heatmaps, box-plots, and correlation plots. Genes were plotted in UCSC Genome Browser [48] and regional association plots in LocusZoom [49]. To access the online results table, visit www.insidegen.com and follow the link ‘Lupus’ then ‘data for scientists’. The table is under title: Expression data associated with different autoimmune diseases.

## Acknowledgements

We thank Dr David L Morris for helpful discussions throughout this work. Philip Tombleson is employed by the Biomedical Research Centre, we thank him for his assistance with data management. The GEUVADIS 1000 Genomes RNA-Seq data was downloaded from the EBI ArrayExpress Portal (accession E-GEUV-1).

## Supporting information

**S1 Table.** SLE GWAS in persons of European Descent (38 loci taken forward for *cis*-eQTL analysis).

**S2 Table.** SLE associated *cis*-eQTL associations deemed to be causal as defined by the JLIM pipeline (this is the output from JLIM).

**S3 Table.** All SLE associated *cis*-eQTL associations by the JLIM pipeline – causal and non-causal associations (provided as a separate XLSX).

**S4 Table.** Functional annotation of SLE candidate genes detected by *cis*-eQTL analysis using RNA-Seq.

**S5 Table.** Number of expression elements that are deemed to have a causal association with the SLE risk SNP.

**S6 Table.** Curated studies of the ImmunoBase Resource.

**S1 Fig.** Overview of the five quantification types used to estimate gene expression using RNA-Seq.

**S2 Fig.** Distribution of joint likelihood *P*-values across RNA-Seq quantification types with 38 SLE GWAS loci.

**S3 Fig.** Specificity of *cis*-eQTLs and candidate genes identified by joint likelihood mapping using SLE GWAS across the five RNA-Seq quantification types.

**S4 Fig.** Regional association plots (+/-250kb) of SLE GWAS in Europeans – showing the nine loci that are causal *cis*-eQTLs and candidate genes from JLIM analysis. The full results of this analysis are in Table 3 of the manuscript and the summary results from the GWAS as provided in S1 Table. Candidate genes are highlighted in red.

**S5 Fig.** SLE associated SNP rs3768792 is a causal *cis*-eQTL for *IKZF2* for a single exon and a single intron.

**S6 Fig.** SLE associated SNP rs7444 is a causal *cis*-eQTL for *UBE2L3* for a single transcript and a single exon.

**S7 Fig.** SLE associated SNP rs9872955 is a causal *cis*-eQTL for *LYST* for a single junction.

**S8 Fig.** Exon and intron numbers for *TYK2* (corresponding to Figure 2). The transcription start site is on the right of the diagram.

**S9 Fig.** Processing of genotype data and principle component analysis. Genotype data in VCF format of 1000Genomes individuals were downloaded from E-GEUV1 (ArrayExpress). Insertion-deletion sites were removed, and bi-allelic SNPs kept only. SNPs with HWE < 0.0001 were removed and the VCF converted to 0,1,2 format using PLINK. Principle component analysis was performed on genotype data using the R package SNPRelate on chromosome 20. The first 3 components were included in the eQTL regression model as well as the binary imputation status (see methods).

**S10 Fig:** Overview of integrative *cis*-eQTL analysis pipeline using 20 autoimmune diseases

